# Inferring condition in wild mammals: body condition indices confer no benefit over measuring body mass across ecological contexts

**DOI:** 10.1101/2023.01.31.524791

**Authors:** Andrea E. Wishart, Adriana L. Guerrero-Chacón, Rebecca Smith, Deborah M. Hawkshaw, Andrew G. McAdam, Ben Dantzer, Stan Boutin, Jeffrey E. Lane

## Abstract

Many studies assume that it is beneficial for individuals of a species to be heavier, or have a higher body condition index (BCI), without accounting for the physiological relevance of variation in the composition of different body tissues. We hypothesized that the relationship between BCI and masses of physiologically important tissues (fat and lean) would be conditional on annual patterns of energy acquisition and expenditure. We studied three species with contrasting ecologies in their respective natural ranges: an obligate hibernator (Columbian ground squirrel, *Urocitellus columbianus*), a facultative hibernator (black-tailed prairie dog, *Cynomys ludovicianus*), and a food-caching non-hibernator (North American red squirrel, *Tamiasciurus hudsonicus*). We measured fat and lean mass in adults of both sexes using quantitative magnetic resonance (QMR). We measured body mass and two measures of skeletal structure (zygomatic width and right hind foot length) to develop sex- and species-specific BCIs, and tested the utility of BCI to predict body composition in each species. Body condition indices were more consistently, and more strongly correlated, with lean mass than fat mass. The indices were most positively correlated with fat when fat was expected to be very high (pre-hibernation prairie dogs). In all cases, however, BCI was never better than body mass alone in predicting fat or lean mass. While the accuracy of BCI in estimating fat varied across the natural histories and annual energetic patterns of the species considered, measuring body mass alone was as effective, or superior in capturing sufficient variation in fat and lean in most cases.

## Introduction

Body mass is among the most frequently measured physical traits of organisms, because it often accounts for much of the observed variation in other traits of interest of the organism (e.g., fecundity; Myers and Master 1983) as well as ecological processes (e.g., trophic structures; Woodward et al. 2005). However, the mechanisms underlying these associations often may depend on the different tissue components comprising body mass, rather than mass itself. For processes in which energy stores or reserves (see Lindström and Piersma 1993 for distinction) are the currency of interest, the non-structural components of body mass that represent metabolizable tissues (fat and lean mass; Krebs and Singleton 1993) are likely more relevant than size or mass alone. For example, relatively high fat stores facilitate successful migration (Bairlein 2002) and hibernation (Humphries et al. 2003), while a higher proportion of lean mass can enhance athletic performance (e.g., takeoff velocity in domestic cats; Harris and Steudel 2002). Understanding relationships between mass and the components of body mass may thus provide greater insight into the mechanisms linking body mass to performance.

A given body mass may be distributed across structural elements of different sizes. Body condition indices (BCIs) attempt to control for this by considering total body mass relative to structural size (e.g., a linear skeletal measurement such as body length, tarsus length, foot length, or a combination thereof; Jakob et al. 1996; Green 2001; Schulte-Hostedde et al. 2001), often to give some indication of an individual’s total lipid stores as a proxy for ‘condition’. Typically, individuals with relatively higher body mass:skeletal size ratios are considered to be in ‘better’ condition than individuals with lower ratios, as the former are assumed to have more energetic stores than individuals of similar structural size, but with a lower metabolizable fraction (Schulte-Hostedde et al. 2001). Accurate measures of ‘condition’ in this energetic sense are widely applicable to scenarios such as livestock breeding programs (e.g., maximizing meat or offspring production), conservation biology (e.g., measuring responses to habitat degradation; Stevenson and Woods 2006; Wikelski and Cooke 2006), and in exploring evolutionary and ecological patterns (e.g., condition-dependent dispersal; Bonte and De La Peña 2009 and migration; Andersen et al. 2000). However, because fat and lean mass have very different energetic storage values (39.6 kJ/g for dry fat, in contrast to 18.1 kJ/g, for dry lean mass; McGilvery 1983), and different functions, metrics that collapse fat and lean into one energetically related fraction can obscure mechanistic relationships between physiology and performance outcomes. Because metabolic rate scales with body size to a lower power than does fat mass, controlling for body size in energetic studies is a logical solution; however, these relationships can vary substantially within and between populations (Glazier 2005). Applications of BCI therefore require a closer consideration of what physiological component of ‘condition’ BCIs are describing (e.g., lean versus fat mass; Schulte-Hostedde et al. 2001).

The dynamic nature of body condition across time (Krebs and Singleton 1993) demands that its correlates and consequences be considered within the context of a species’ natural history (Molnár et al. 2009). The composition of a body in ‘good condition’ in an adaptive sense (i.e., that which is associated with increased survival and/or fitness, *sensu* Wilson and Nussey 2010), should therefore be expected to vary over time depending on circannual energetics. Fat reserves are an important source of metabolizable energy (Jenni and Jenni-Eiermann 1998) that fuel organisms through energetically-expensive behaviours or during periods of energetic shortfall. For example, individuals with higher fat fractions entering hibernation are more likely to survive over winter and breed successfully the following spring (Boyer and Barnes 1999). Fatter is not always better though: during the active season, carrying more body mass can decrease running speed (Trombulak 1989) and alter circulating hormones (Taylor et al. 1982). Arboreal species may trade off fat stores for locomotion (Dittus 2013), and for species that primarily store energy off-body as external food caches, fat reserves may not confer the same advantages during resource-scarce seasons as they may for species that store energy exclusively on-body. Furthermore, a higher lean fraction may be associated with increased activity, as shown in captive rats (*Rattus norvegicus*; Swallow et al. 2010). The body composition associated with better performance is therefore likely to be highly dependent on natural history, as well as on the various activities an animal undertakes throughout the year (Wells et al. 2019).

Body mass and structural size can be measured from live animals, giving BCIs an advantage over chemical composition analyses that require lethal sampling (Reynolds and Kunz 2001), especially in studies where repeated measures within individuals are valuable. In many species, BCIs correlate well with chemical quantification of fat mass in vertebrates including birds (Chang and Wiebe 2016) and reptiles (Weatherhead and Brown 1996), and invertebrates including arthropods (Jakob et al. 1996; Moya-Laraño et al. 2008; Kelly et al. 2014). However, BCIs are not without drawbacks. Acquiring the structural size component(s) of the index can be challenging. Krebs and Singleton (1993) warned of low repeatability attributable to measurement bias across different observers (although this can be greatly reduced by taking replicate measurements; Blackwell et al. 2006). While estimating lengths of long bones yields accurate estimates when measured from museum specimens (e.g., Dobson 1992), long bones can be challenging to measure on live, unanesthetized animals in field conditions (Green 2001). Even under anesthesia, measurement error can persist (e.g., Martin et al. 2013). Because skeletal morphology often reflects natural history, as selection favours certain shapes for certain lifestyles (e.g., arboreal vs. fossorial, e.g. Shimer 1903), the body component of interest may not be accurately reflected by BCI if the linear measurement(s) of size does not correlate well with overall body size. Furthermore, although BCI can correlate more strongly with lean mass rather than fat mass (Schulte-Hostedde et al. 2001), the efficacy of BCI in predicting the masses of body components may be no more effective than prediction through body mass alone (e.g., body mass predicts fat stores in bats just as well as BCI; McGuire et al. 2018, Wells et al. 2019). Finally, since BCI estimates the non-skeletal component of body composition in a general sense, its interpretation and therefore relevance may differ depending on the energetic aspects of the natural history of the focal species (e.g., for a fat-storing hibernator versus a food-caching non-hibernator). We thus expect the correlation between BCIs and body composition (i.e., fat and lean mass) to vary both across species and timing in annual cycles, reflecting the dynamic nature of energy budgets in seasonally dependent activities.

We studied wild populations of three different mammal species to evaluate concordance between BCIs and the mass of fat and lean tissue, and determined whether these relationships differ according to the species’ primary energy storage mode (e.g., on-body fat storage vs. off-body food caching). We hypothesized that the relationship between BCI and body composition depends on both the extent to which the species relies on on-body energy stores (i.e., fat) for overwinter survival, and with the expected energetic balance within a stage of the annual cycle (i.e., season).

We selected three species in the family Sciuridae which differ in patterns of energy storage and metabolic demands: North American red squirrels (*Tamiasciurus hudsonicus*, hereafter, red squirrels), black-tailed prairie dogs (*Cynomys ludovicianus*, hereafter prairie dogs), and Columbian ground squirrels (*Urocitellus columbianus*, hereafter ground squirrels). We also compare ground squirrels before and after hibernation to characterize within-species shifts in energy stores. Red squirrels harvest and cache conifer cones in a central larder (‘midden’; Smith 1968) from late summer through autumn (Fletcher et al. 2010) and weigh ~230-250 g on average as adults (Boutin and Larsen 1993). Red squirrels remain euthermic throughout winter without using torpor (Brigham and Geiser 2012), but are able to maintain low levels of energy expenditure through behavioural adjustments (Humphries et al. 2005) and the use of well insulated nests (Studd et al. 2016). They are not known to gain significant amounts of fat prior to winter, during which they rely on cached resources for energy (Wishart 2023).

Prairie dogs can weigh up to 1710 g, but show high within- and between-individual variation in body mass (Kusch et al. 2021). Throughout most of their range, prairie dogs are active throughout the winter, however they are capable of hibernation (Lehmer et al. 2006). In southwestern Saskatchewan, where our study population is located at the northern edge of the species distribution, they hibernate for ~4 months during winter (Gummer 2005). Prairie dogs of both sexes increase overall body mass, and fat mass specifically, leading up to winter (Lehmer and Van Horne 2001; Kusch et al. 2021).

Ground squirrel body mass varies substantially across their active season (~400 g at emergence from hibernation in spring, to up to ~700 g prior to immergence in late summer; (Boag and Murie 1981, Dobson et al. 1992). These obligate hibernators are notable for their short active season (~4 months) and extended time spent in hibernation (~8 months) each year (Dobson et al. 1992). Ground squirrels reach lower minimum body temperatures (0 °C) and have longer torpor bout durations (~390 hours for males; Young 1990) than prairie dogs. We 1) characterized morphology and body composition of these three species in context of their seasonal energetic demands, 2) evaluated the correlation of BCI with body composition variables (lean and fat mass), and 3) determined whether BCI confers any benefit relative to body mass in predicting body composition variables (i.e., whether the correlation between BCI fat or lean mass was higher than between body mass fat or lean mass. We expected hibernators (ground squirrels and prairie dogs) to have the highest fat fraction during the pre-winter season, given the importance of on-body energetic reserves to sustain hibernation, compared to non-hibernating red squirrels which rely mostly on hoarded food, and compared to post-emergence ground squirrels in spring who have metabolized fat stores overwinter (Fletcher et al. 2013). Because fat stores are expected to represent the bulk of the pre-hibernation weight gain in the lead up to winter, we also expected high concordance between BCI and fat mass in pre-winter hibernators. Conversely, we expected the lowest concordance between BCI and fat mass to be in red squirrels, as lean mass is likely to be more important to sustained caching activity (Fletcher et al. 2015).

## Materials and Methods

### Study sites and population monitoring

We sampled free-ranging, non-breeding adults from populations within the northern regions of their respective ranges in Canada: red squirrels in the southwest Yukon (61° N, 138° W, ~ 850 m above sea level (a.s.l.)), prairie dogs in Grasslands National Park, Saskatchewan (49° N, 107° W, ~770 m a.s.l.), and ground squirrels in Sheep River Provincial Park, Alberta (50° N, 114° W, ~ 1500 m a.s.l.). For all populations, we collected data through live-trapping. All populations had been monitored for at least one year prior to data collection for the present study. For detailed descriptions of population and reproductive monitoring, see McAdam et al. (2007) and Dantzer et al. (2020) for red squirrels, Kusch et al. (2020) for prairie dogs, and Lane et al. (2019) for ground squirrels. Briefly, all individuals received permanent uniquely marked ear tags (National Band and Tag Company, Newport, KY) upon first trapping. We included only adults (individuals one year of age or older) to minimize effects of skeletal growth dynamics. For ground squirrels and prairie dogs, we knew ages for all individuals captured at first emergence following birth. For red squirrels and some older ground squirrels and prairie dogs, exact ages were not known, but we could confidently remove young-of-the-year based on size and breeding/nipple status on first trapping (red squirrels excluded if under 150 g, ground squirrels excluded if under 400 g, prairie dogs excluded if under 800 g; for all species, females also excluded if they had small pink nipples at first trapping). We excluded all pregnant (assessed by abdominal palpations) and lactating (assessed by milk expression) females to remove variance related to maternal investment in offspring. Sample sizes for each sex/species/season group are summarized in Table 1.

**Table 1.**
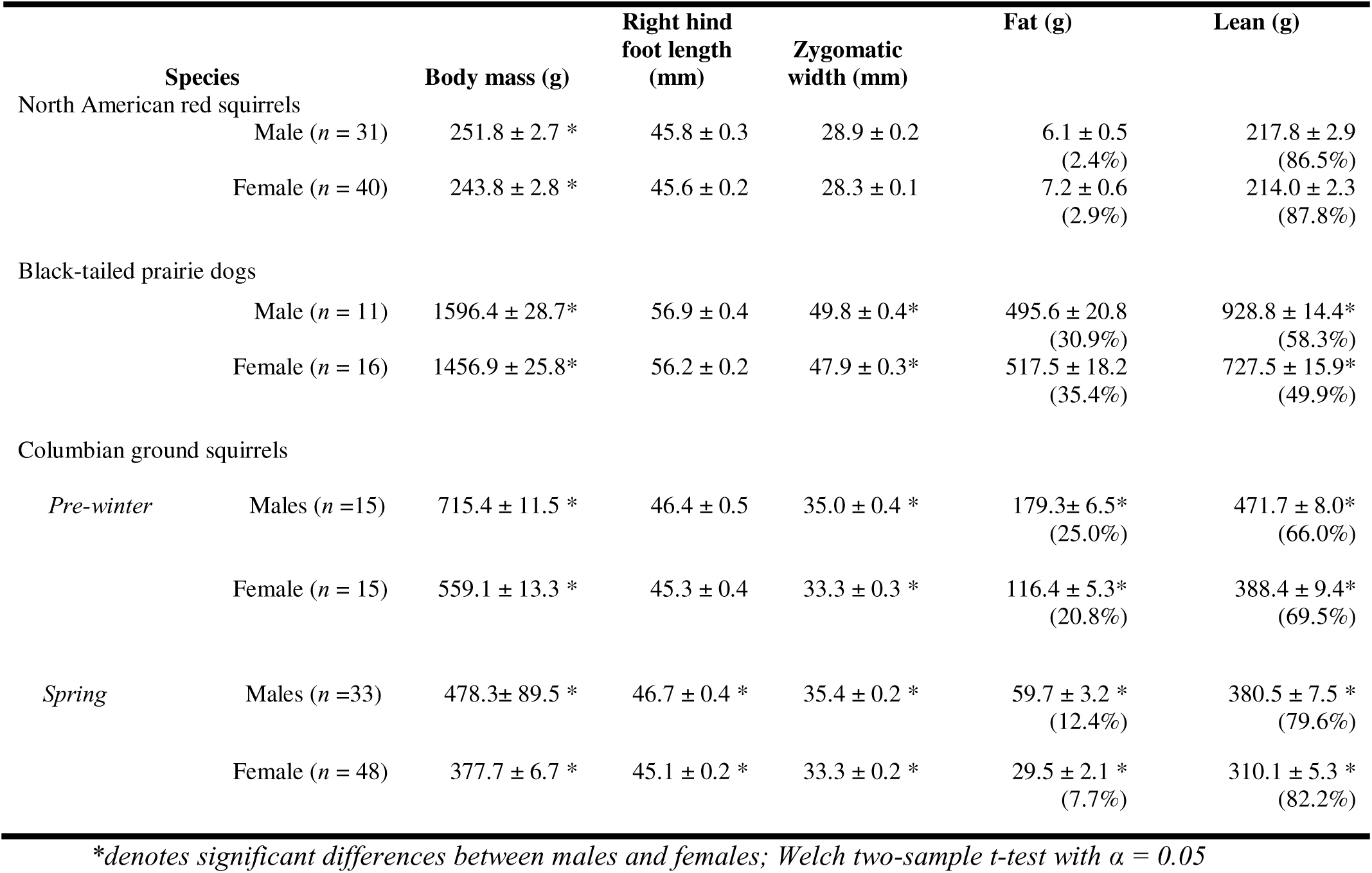
Summary of morphometric data (body mass, right hind foot length, and zygomatic width) and body composition (fat mass, lean mass) for adult non-breeding male and female North American red squirrels (pre-winter), black-tailed prairie dogs (pre-winter), and Columbian ground squirrels (pre-winter and spring). Values reported as mean ± standard error of the mean (SEM). Sample sizes for each sex (♂ = males, ♀ = females) are indicated in the species column.

| Species |  | Body mass (g) | Right hind foot length (mm) | Zygomatic width (mm) | Fat (g) | Lean (g) |
| --- | --- | --- | --- | --- | --- | --- |
| North American red squirrels |  |  |  |  |  |  |
| | Male ($n = 31$) | 251.8 ± 2.7 * | 45.8 ± 0.3 | 28.9 ± 0.2 | 6.1 ± 0.5<br>(2.4%) | 217.8 ± 2.9<br>(86.5%) |
| | Female ($n = 40$) | 243.8 ± 2.8 * | 45.6 ± 0.2 | 28.3 ± 0.1 | 7.2 ± 0.6<br>(2.9%) | 214.0 ± 2.3<br>(87.8%) |
| Black-tailed prairie dogs |  |  |  |  |  |  |
| | Male ($n = 11$) | 1596.4 ± 28.7* | 56.9 ± 0.4 | 49.8 ± 0.4* | 495.6 ± 20.8<br>(30.9%) | 928.8 ± 14.4*<br>(58.3%) |
| | Female ($n = 16$) | 1456.9 ± 25.8* | 56.2 ± 0.2 | 47.9 ± 0.3* | 517.5 ± 18.2<br>(35.4%) | 727.5 ± 15.9*<br>(49.9%) |
| Columbian ground squirrels |  |  |  |  |  |  |
| *Pre-winter* | Males ($n = 15$) | 715.4 ± 11.5 * | 46.4 ± 0.5 | 35.0 ± 0.4 * | 179.3 ± 6.5*<br>(25.0%) | 471.7 ± 8.0*<br>(66.0%) |
| | Female ($n = 15$) | 559.1 ± 13.3 * | 45.3 ± 0.4 | 33.3 ± 0.3 * | 116.4 ± 5.3*<br>(20.8%) | 388.4 ± 9.4*<br>(69.5%) |
| *Spring* | Males ($n = 33$) | 478.3 ± 89.5 * | 46.7 ± 0.4 * | 35.4 ± 0.2 * | 59.7 ± 3.2 *<br>(12.4%) | 380.5 ± 7.5 *<br>(79.6%) |
| | Female ($n = 48$) | 377.7 ± 6.7 * | 45.1 ± 0.2 * | 33.3 ± 0.2 * | 29.5 ± 2.1 *<br>(7.7%) | 310.1 ± 5.3 *<br>(82.2%) |

\*denotes significant differences between males and females; Welch two-sample $t$ -test with $\alpha = 0.05$

### Morphometric measurements

We measured body mass and size for all live-trapped individuals (Table 1). We weighed each prairie dog to the nearest 5 g using a Pesola spring scale (Pesola AG, Baar, Switzerland), and weighed red squirrels and ground squirrels to the nearest 1 g on an electronic balance (Ohaus C5 Series 2000 g). We measured zygomatic arch width (‘ZW’) to the nearest millimeter using calipers (analogue for red squirrels and prairie dogs; digital for ground squirrels). We measured right hind foot (‘RHF’) length from heel to longest toe (excluding claw) to the nearest millimeter using a ruler fit with a heelstop at 0 mm. We measured both ZW and RHF three times per handling, and used the mean value for analyses.

We took these measurements during the same handling occurrence as body composition analyses for all red squirrels, all prairie dogs except one, and most ground squirrels. For the remaining prairie dog and ground squirrels, we used skeletal measurements taken on the nearest date to when composition analyses were completed. The prairie dog was an adult and unlikely to be growing, so we used measurements taken 81 days prior. The median interval between skeletal measurements and composition scans for ground squirrels was 0 days (range: 0-120). We excluded data from ground squirrels that were younger than 3 years of age if their skeletal measurements were taken more than two weeks before or after the date of body composition and mass measurements, as younger squirrels may still be growing structurally (Dobson 1992). A single yearling (female) remained in the spring dataset, so we removed all yearling ground squirrels.

### Body composition

We measured pre-winter body composition of red squirrels between late-September and mid-October in 2018 and 2019, prairie dogs in late October 2018 (due to variable torpor patterns in prairie dogs, exact immergence date into hibernation was not known), and ground squirrels between late-July and late-August 2019 within a week of immergence for most individuals (median = 3.5 days pre-immergence; range = 0-15). We measured spring body composition of ground squirrels for which we were confident were captured within a few days of emergence from hibernation (median = 1 day post-emergence; range = 0-3) between mid-April and early May 2019. Most individual ground squirrels were measured in one season only, but 21 individuals were measured in both seasons. As we do not expect ground squirrels to change skeletal size substantially after they have reached maturity, we recorded morphometric data once per individual.

We used a quantitative magnetic resonance (QMR) body composition analyzer (EchoMRI-1600, Echo Medical Systems, Houston, TX) to measure absolute lean and fat mass (g). This technology provides a relatively non-invasive method of estimating body composition precisely, and accurately (Tinsley et al. 2004). The remaining components not captured as fat and lean mass are free water and skeletal mass (McGuire and Guglielmo 2010). Quantities measured using QMR correlate well with carcass-derived quantities for numerous species across multiple taxa, most relevant here being rodents, including laboratory rats (*Rattus norvegicus domestica*; Johnson et al. 2009) and house mice (*Mus musculus domesticus*; Jones et al. 2009). The QMR approach allows for repeated measures of live animals, both awake and sedated (Tinsley et al. 2004; McGuire and Guglielmo 2010; Zanghi et al. 2013a, b).

We housed our QMR system in a custom-designed trailer to enable transportation to each study site. The trailer was climate-controlled to stabilize the temperature, as the magnet within the QMR system is temperature-sensitive. We targeted an ambient temperature of 21 °C as per manufacturer recommendations, although field conditions widened the range of stabilized temperatures to ± 7 °C. We calibrated the system daily to a 943 g canola oil standard at the stabilized temperature, which was held for at least five hours prior to scanning animals (following Guglielmo et al. 2011). Our QMR system was outfitted with an additional antenna to measure animals from 100 g up to 1600 g to accommodate the range of body masses of the three species. Details of similar systems are described elsewhere (McGuire and Guglielmo 2010), and we followed similar protocols here. We live-trapped squirrels in the field, and transported them to the trailer. We placed each squirrel in a clear plexiglass holding tube with perforations to allow ample airflow to the animal, then inserted the tube into the QMR chamber. We recorded body composition through a minimum of two scans, reporting the average values for each individual. In 2019, we administered a mild sedative via thigh intramuscular injection to red squirrels prior to scanning and collecting morphometric data (100 µg/kg of dexmedetomidine, reversed by 1 mg/kg atipamezole) to minimize stress and movement during scans. Sedation was not necessary for the semi-fossorial prairie dogs or ground squirrels, who remained still, and even fell asleep in the chamber.

### Calculating body condition indices (BCIs)

We retained individuals in the dataset for which we had measurements for all of the following: ZW, RHF, body mass, and body composition. Within each species, we calculated coefficients of variation for all variables, and analyzed relationships among variables. While principal components analysis has been used previously as a general measure of structural size to derive BCI (Schulte-Hostedde et al. 2005), we determined that it was not appropriate for our dataset because correlation coefficients between ZW and RHF were not always positive (Supplementary Fig. S1). Instead, we selected the single skeletal measure that had the greatest coefficient of variation (CV) to generate a residual index (Jakob et al. 1996). Red squirrels showed a negative, but non-significant, relationship between RHF and ZW (Supplementary Table S1) so we chose to use ZW to generate the BCI for this species (‘ZW index’). The relationships between RHF and ZW appeared to differ in direction between male and female prairie dogs; however, neither correlation was significant, so we generated a ZW index for this species. In ground squirrels, only data for females pre-winter showed a significant (positive) relationship. Females in spring, and males in both seasons, showed no significant relationship between RHF and ZW. We therefore used RHF to generate the single-metric BCI (‘RHF index’) for ground squirrels.

To calculate each BCI, we regressed either RHF (log transformed and standardized to mean of zero and unit variance) alone on body mass (log transformed and standardized as above; generating the RHF index), or ZW (log transformed and standardized as above) alone on body mass (log transformed and standardized as above; generating the ZW index). Residual plots are shown in Supplementary Fig. S2. We calculated BCIs within season (pre-winter or spring) and within sex for each species using separate regressions. Because some CV values within species/seasons were similar for RHF and ZW, we also ran models using the BCI derived from the alternate skeletal measurement (Supplementary Online Material). We indicate in the results when results differed from the primary BCI model.

In addition to the residual-derived BCIs described above, we also calculated the scaled mass index (SMI; Peig and Green 2009) because it accounts for scaling relationships of linear measurements (e.g., hind foot length), tissue components, and overall body size. We then ran parallel statistical analyses as described below, substituting SMI values for BCI values in all models (see Supplementary Table S11). We note in the Results when the relationships differed between SMI and BCI.

### Statistical analyses

We performed all analyses in R (v.4.0.3, R Core Team (2020)). We compared morphological measures between the sexes within each species (and within season for ground squirrels) using a Welch’s two-sample t-test. To assess the efficacy of the selected BCI for each species in predicting body composition variables, we modeled, separately for each species (and for ground squirrels, separately for each season), linear models for fat mass (g) and lean mass (g), each predicted by the selected BCI interacting with sex. We also modeled fat and lean mass predicted by body mass interacting with sex. We compared the fit of the BCI model and body mass model for each component for each species using Akaike’s information criterion adjusted for small sample size (Burnham and Anderson 2002), confirming no missing data. We considered the model with the lowest AIC to be the best fit for explaining the variation in the response variable, among candidate models. Models that were different by >2 AICc were considered to be different from one another. To compare the utility of BCI to predict fat and lean mass across species, we fit a linear model for data from all three species (including both seasons for ground squirrels) together. For fat and lean mass separately, we defined an interaction term between species-specific BCI and species as the independent variable. We fit a similar model set with body mass (scaled within species) instead of BCI.

## Results

### Morphological measurements

Males were significantly heavier than females in all three species: males were 3.3% heavier in red squirrels, 9.6% heavier in prairie dogs, 27.9% heavier in pre-winter ground squirrels, and 26.6% heavier in spring ground squirrels (Table 1). Right hind foot length was different between sexes only for ground squirrels in spring, with male RHF 1.6 mm longer than in females. Zygomatic width was significantly larger in male prairie dogs and ground squirrels than females, but similar between red squirrel males and females.

### Body composition

In autumn, red squirrels showed the lowest percent body fat, with 2.4% in males and 2.9% in females (Table 1) while prairie dogs had the highest percent body fat, with 30.9% and 35.4% in males and females, respectively. Ground squirrels were the only species to have statistically significant sex differences in fat (in both seasons). They showed their highest body fat levels pre-winter before entering hibernation (25.0% for males, 20.8% for females, t = −7.48, d.f. = 26.9, *p* < 0.001), and lowest in spring (12.4% fat for males, 7.7% fat for females, t = −7.97, d.f. = 58.5, *p* < 0.001).

### Relationship between BCI, body mass, and body composition

The relationships between BCI/body mass and fat/lean mass was positive in almost all cases (Figures 1–4; the exception being for male prairie dog BCI and lean mass, Figure 2b; Supplementary Tables S3-6). For red squirrels, the BCI and body mass models performed similarly in predicting fat, but correlations with fat were weak (adjusted *R*^2^ < 0.22) for both BCI and mass (Supplementary Table S3, Figure 1). Red squirrel lean mass was better predicted by both BCI and mass models than fat mass, with the body mass model providing a better fit. This pattern held for the alternate RHF BCI models (Supplementary Table S7). For prairie dogs, both ZW BCI and body mass predicted fat well, but the body mass model was a better fit for both fat and lean mass (Supplementary Table S4, Figure 2). For pre-winter ground squirrels, the two models for fat were indistinguishable based on AICc and had similar correlation strengths (Supplementary Table S5, Figure 3). The body mass model was a better fit for lean mass. This pattern held for spring, with BCI and body mass models being similar for fat, and the body mass model providing a better fit for lean mass (Supplementary Table S6, Figure 4). Patterns did not change when using ZW BCI instead (Supplementary Tables S7-8). Additionally, for ground squirrels, the variance in fat explained by the BCI and body mass models was higher pre-winter (adjusted *R^2^* = 0.88) compared to the spring (adjusted *R^2^* = 0.69). The only comparison in which the pattern for scaled mass index (SMI) differed from the BCI model sets was for red squirrel fat: this was also the only instance when the SMI model was a better fit than the body mass model (adjusted *R^2^* = 0.25; Supplementary Table S11). Otherwise, the body mass model outperformed the SMI model in all parallel analyses.

**Figure 1.**
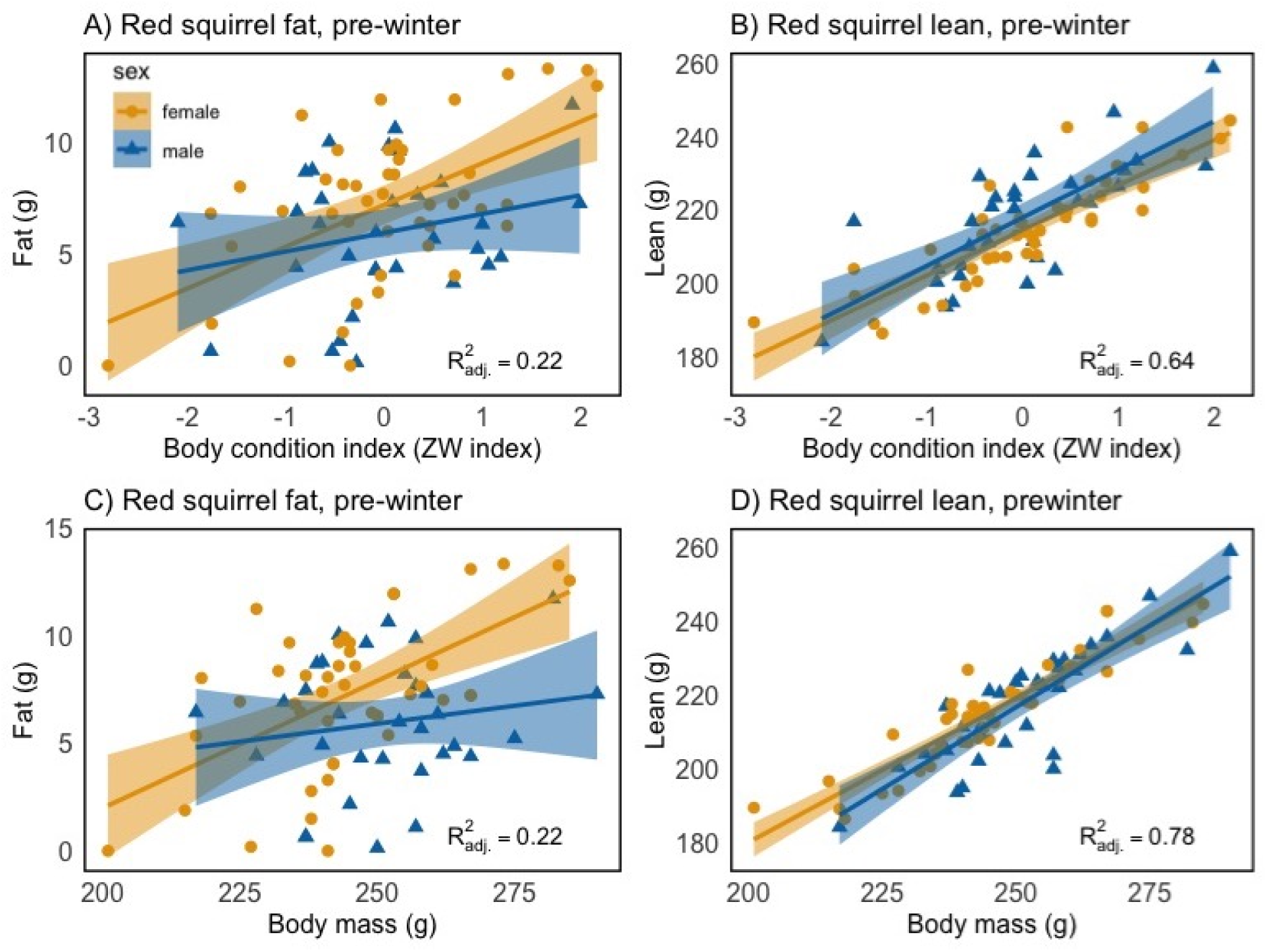
Body composition (fat mass [A,C] and lean mass [B,D] in grams) as a function of zygomatic-derived body condition index (ZW index, A-B) and body mass (C-D) for male (blue triangles) and female (orange circles) North American red squirrels pre-winter.

**Figure 2.**
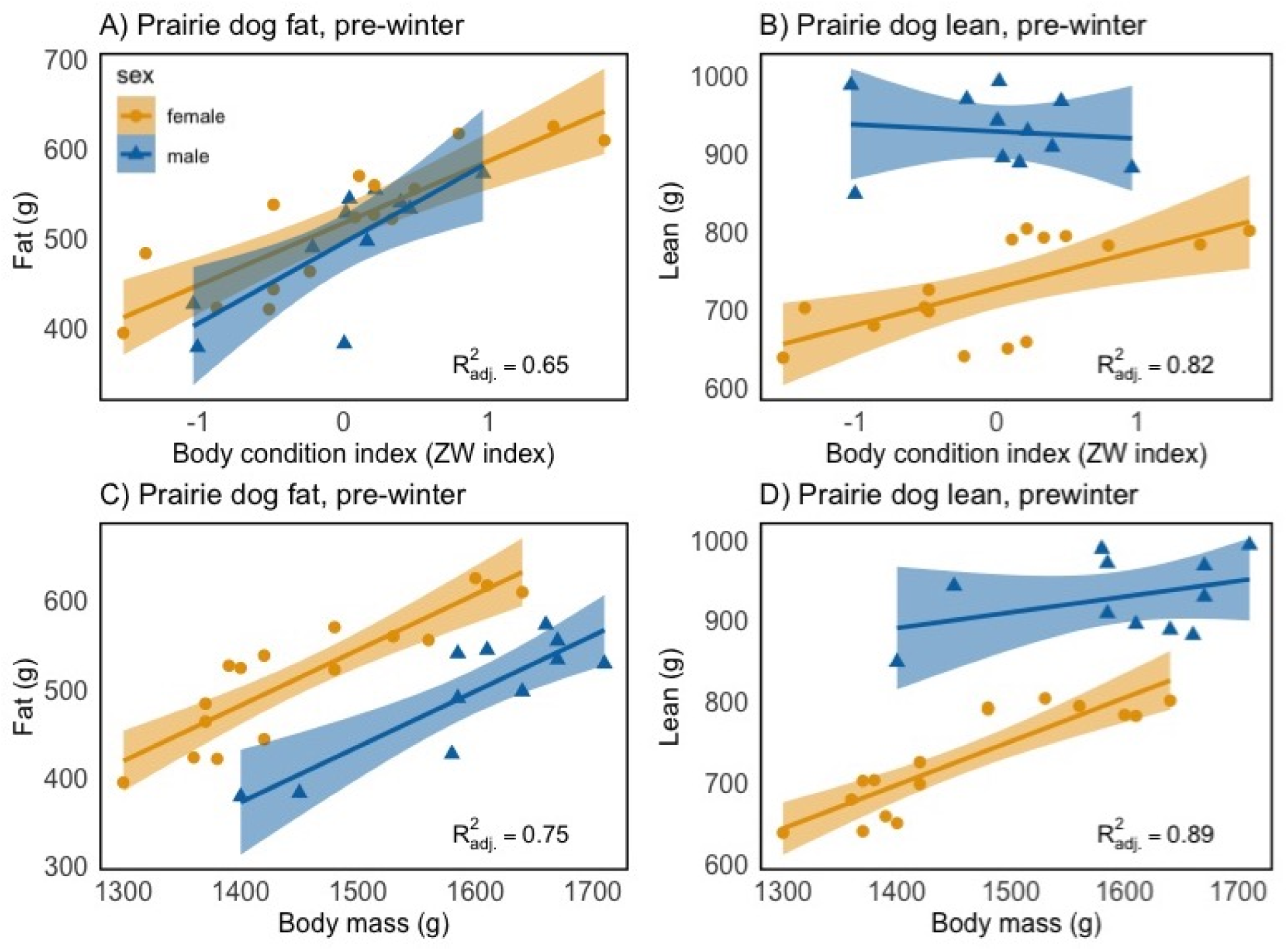
Body composition (fat mass [A,C] and lean mass [B,D] in grams) as a function of zygomatic-derived body condition index (ZW index, A-B) and body mass (C-D) for male (blue triangles) and female (orange circles) adult non-breeding black-tailed prairie dogs pre-winter.

**Figure 3.**
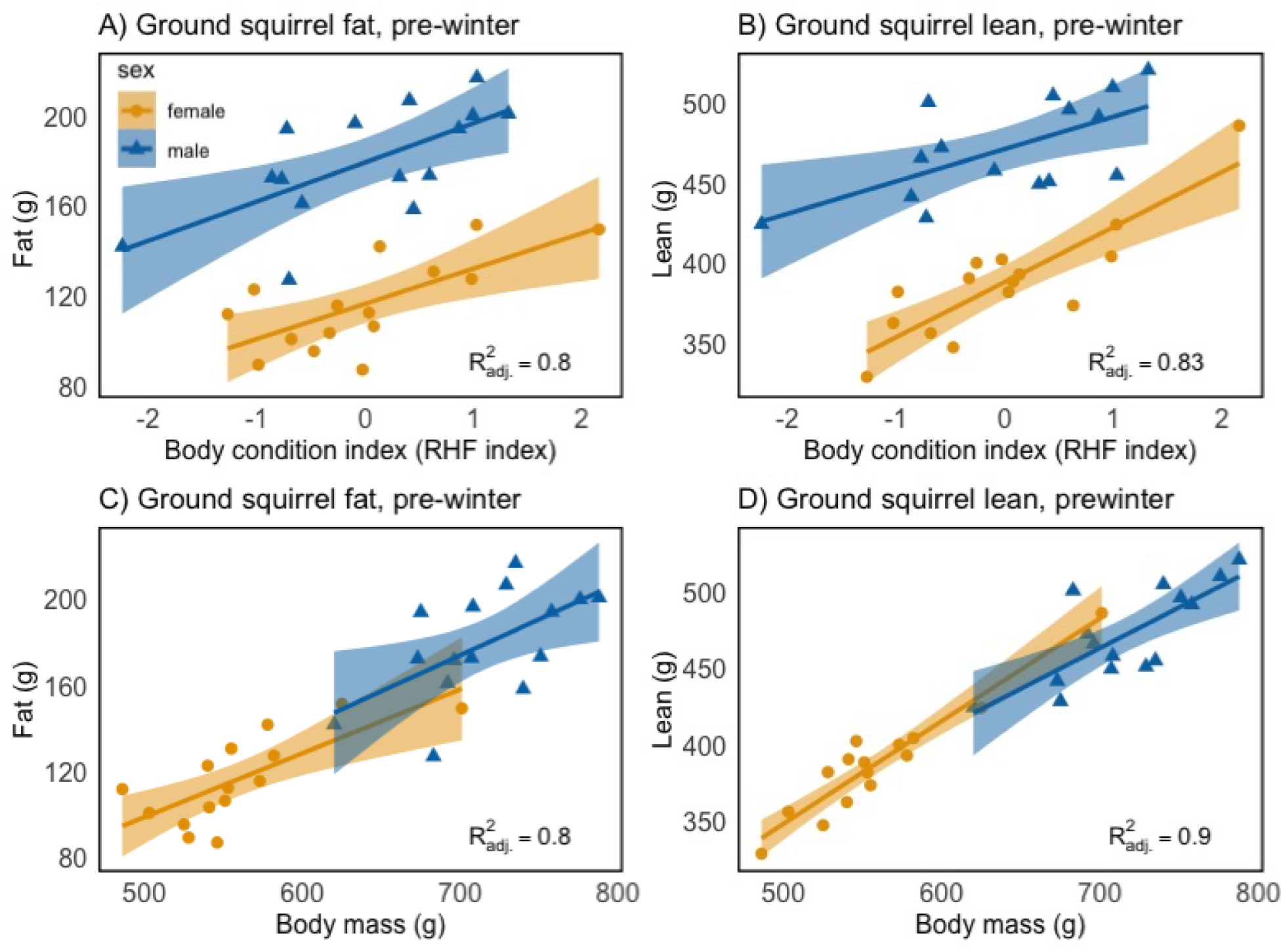
Body composition (fat mass [A,C] and lean mass [B,D] in grams) as a function of right hind foot-derived body condition index (RHF index, A-B) and body mass (C-D) for male (blue triangles) and female (orange circles) adult non-breeding Columbian ground squirrels pre-winter.

**Figure 4.**
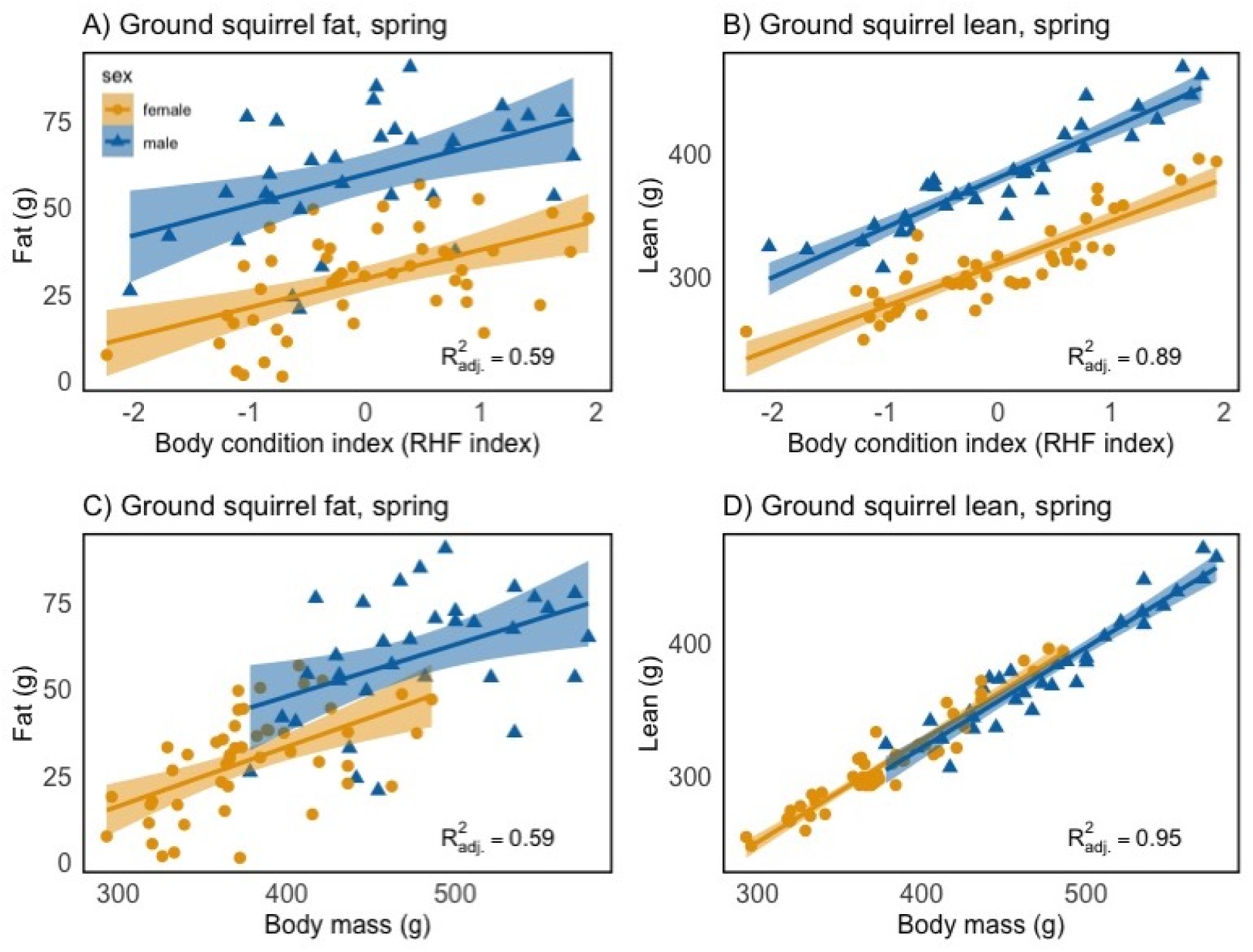
Body composition (fat mass [A,C] and lean mass [B,D] in grams) as a function of right hind foot-derived body condition index (RHF index, A-B) and body mass (C-D) for male (blue triangles) and female (orange circles) adult non-breeding Columbian ground squirrels in spring.

In the models including data from all species (and for ground squirrels, both seasons), the strength of the correlations in both BCI and body mass model sets were strong (adjusted *R^2^* > 0.9 for all) and similar within each component. However, body mass models were consistently a better fit than the BCI models (ΔAICc = 26.7 for fat, 18 for lean mass, Figure 5, Supplementary Table S12).

**Figure 5.**
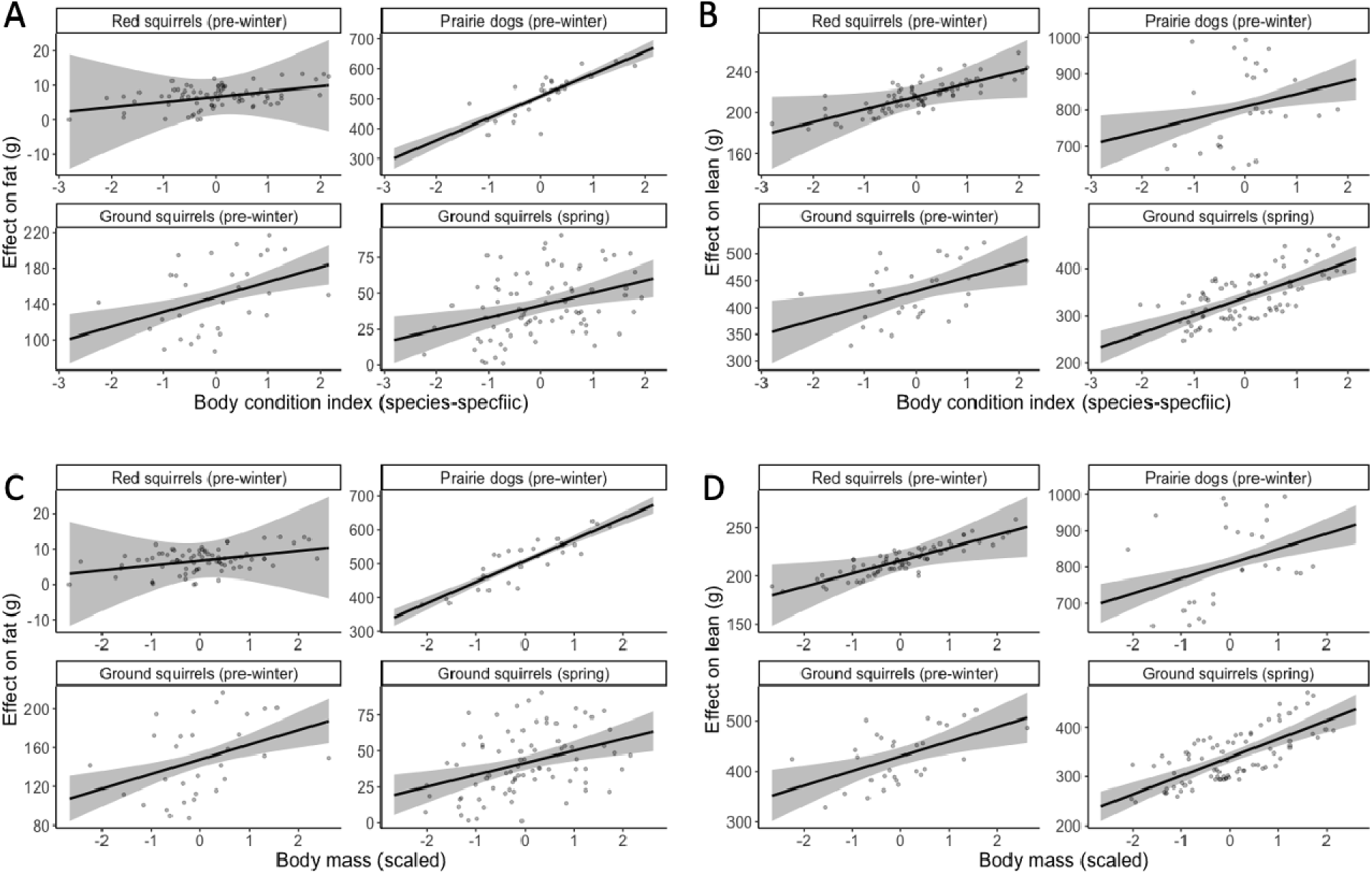
Partial plots showing the relationship between (A, B) species-specific body condition indices and (C, D) body mass on (A, C) fat and (B, D) lean mass in two linear models for adult non-breeding North American red squirrels (pre-winter), black-tailed prairie dogs (pre-winter), and Columbian ground squirrels (pre-winter and spring).

## Discussion

We demonstrate that the utility of body condition indices in predicting physiologically relevant tissues, specifically fat, is conditional on the expected energetic state of a species given ecological and natural history considerations. If BCI is a reliable indicator of ‘condition’ as it relates to on-body energy stores, it should be strongly correlated with fat mass. However, when individuals were expected to be in a leaner state (e.g., food-caching red squirrels, spring ground squirrels), correlations between BCI and fat were low to moderate. When individuals were expected to have higher fat stores (e.g., pre-hibernation prairie dogs and ground squirrels), BCI models predicted fat well, following similar patterns in seabirds (Jacobs et al. 2012). Furthermore, the relationship between BCI and fat varied among species (and season, for ground squirrels). This contingency creates a problem for using BCI to infer size of masses of fat; specifically, the accuracy of the tool differs along the gradient of variation it is being used to infer. In nearly all cases, BCI was positively correlated with lean mass. Because fat and lean mass have significantly different energetic values (McGilvery 1983, Jenni and Jenni-Eiermann 1998), interpretations of BCI as they relate to metabolizable energy should take into account species-specific natural history and annual energetic patterns. For example, some red squirrels that would have ranked as lower condition based on BCI had nearly twice as much fat as some individuals that had higher BCI values. Furthermore, models fit with body mass were almost always more highly correlated with fat and lean mass than models with BCI, both within and across species.

The selection of these three species, and the time of year at which they were studied, provides insight into how body composition manifests in BCI when individuals are in a peak positive energy balance after accumulating surplus energy to sustain them through upcoming energetic shortfalls, and when they are expected to have depleted much of that accumulated energy. By investigating these relationships at the ends of the continuum of energetic states that organisms may experience throughout the year, we demonstrate that interpretation of ‘condition’ indices should consider seasonal variation in energetic demands. For example, red squirrels primarily store energy as cached food, so were expected to carry little fat. In comparison, we expected prairie dogs to have high fat stores to sustain them through inefficient hibernation (Gummer 2005; Hawkshaw 2022), and indeed they had the highest percent body fat of all three species. The intermediate ground squirrels were the only species to show sex differences in fat, likely arising from asymmetry in reproductive phenologies between males and females (males arouse from torpor earlier to undergo spermatogenesis, relying on stored energy, while females arouse later and have access to more food in the environment to support the cost of lactation, Broussard et al. 2005). These results illustrate that the relative importance of fat and lean mass is likely to vary with seasonal activities interacting with the natural history of the organisms.

Almost every relationship between either predictor variable (BCI or body mass) and body component (fat or lean mass) was positive, save for male prairie dog lean mass. Echoing Schulte-Hostedde et al. (2001), BCIs are capturing variation in both components; however, we did not find BCIs to discriminate between the independent variation in each component. We found that in general, lean mass had less variation around the line of best fit than fat mass, reflective of previous studies on small-non hibernating mammals that also found that BCIs tend to be more effective in predicting lean dry mass and water as compared to fat mass (Schulte-Hostedde et al. 2001; Tidhar and Speakman 2007). Given the dynamic nature of body composition in hibernating species, inferring fat levels from BCIs can be particularly difficult since many studies that use residual-derived BCIs to assess fat assume that lean mass scales with body size, and that fat mass varies with condition (McGuire et al. 2018).

In some contexts, like large scale comparisons across species, body mass or BCIs could be a rough proxy for both fat and lean mass. However, for comparisons among individuals or within individuals of the same species, or research questions that require the precise estimation of energy derived from the changes in body mass/BCI, a more precise technique like QMR is needed (e.g., to estimate energy drawn from either fat lean mass during hibernation, Mejías et al. 2022; or migration, McGuire et al., 2022; Kelsey and Bairlein, 2019).

We have shown that the utility of body mass and/or BCI to describe energy-relevant components is conditional on natural history and annual energetic cycles. This demands not only within-species validation if BCI is to be used, but also careful consideration of how these relationships may change across seasons or energetic states. If BCIs are only predictive of fat in the upper end of the range of variation in fat stores, the value of BCI in comparing across individuals breaks down, especially across individuals with relatively lower fat masses. As shown within Columbian ground squirrels across two seasons, BCI (and the associated challenges and handling stress involved in obtaining linear measurements of skeletal size can impart on animal subjects) is not uniformly informative, therefore bringing into question its utility. When balancing animal welfare, time, and research budgets with desired outcome of informative data, we advise researchers to consider whether more precise measurements of body composition are needed, or if simply measuring body mass is sufficient to capture the variation of interest.

Ultimately, this study strengthens the case for using body mass as a covariate to capture general variation in fat and lean masses in most scenarios rather than the less informative and more difficult to assess BCI. BCIs rarely conferred an advantage over mass in predicting fat and lean mass across the range of species, seasons, and ecological contexts considered here. Further research into mass and composition dynamics across seasons and in different energetic contexts will help determine the extent to which such relationships hold outside seasons of expected extremes of energy budgets.

## Acknowledgements

We thank Agnes MacDonald for long-term access to her trap-line, and to the Champagne and Aishihik First Nations for allowing us to conduct fieldwork related to red squirrels within their traditional territory. Fieldwork related to prairie dogs was conducted in Treaty 4 territory, the traditional territory of the Oceti Sakowin and Niitsitpiis-stahkoii, and the homeland of the Métis Nation. Fieldwork related to ground squirrels was conducted in Treaty 7 territory, the traditional territories of the Blackfoot Confederacy (Siksika, Piikani, and Kainai First Nations), the Tsuut’ina First Nation, the Stoney Nakoda (Chiniki, Bearspaw, and Wesley First Nations) and homeland of the Métis Nation. We thank the many volunteers, graduate students, and field assistants, including Jillian Kusch, Tessie Aujla, Jack Hendrix, Ashley Mills, Emily Kelvin, Megan Miller, M. Alejandra Hurtado, and Gabriela Heyer for field data collection, and Grasslands National Park staff for assistance with permitting and logistical support. AEW thanks Dr. Kurtis Swekla and Dr. Todd Shury for professional veterinary services in developing anesthesia protocols for red squirrels.

## Declarations

### Ethics approval

All procedures were approved by the University of Saskatchewan Animal Research Ethics Board. Fieldwork was completed under permits issued by Yukon Territorial Government (red squirrels); Grasslands National Park and the Saskatchewan Ministry of the Environment (prairie dogs); and Alberta Parks (ground squirrels).

### Consent to participate

Not applicable.

### Consent for publication

Not applicable.

### Availability of data

Data are archived on FigShare at the following private link: https://figshare.com/s/51b0edc3df7e82bb9609. Data will be made public with an associated DOI upon acceptance.

### Code availability

Code for analysis is available on GitHub at https://github.com/aewishart/bci-composition.

