## Supplementary Fig. S1 for "Inferring condition in wild mammals: body condition indices confer no benefit over measuring body mass across ecological contexts"

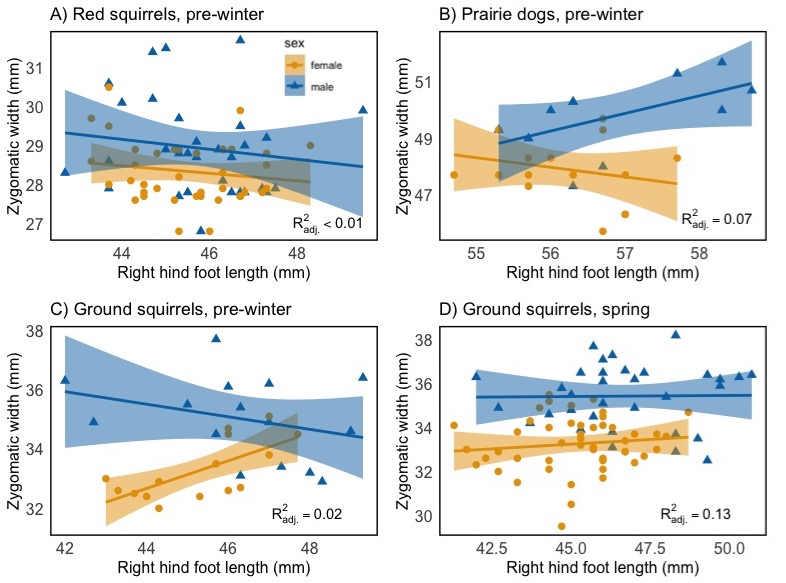

**Supplementary Figure S1.** Correlation between zygomatic width (mm) and right hind foot length (mm) for A) North American red squirrels pre-winter, B) black-tailed prairie dogs pre-winter, and Columbian ground squirrels C) pre-winter and D) in spring (D) for both males (blue triangles) and females (yellow circles).

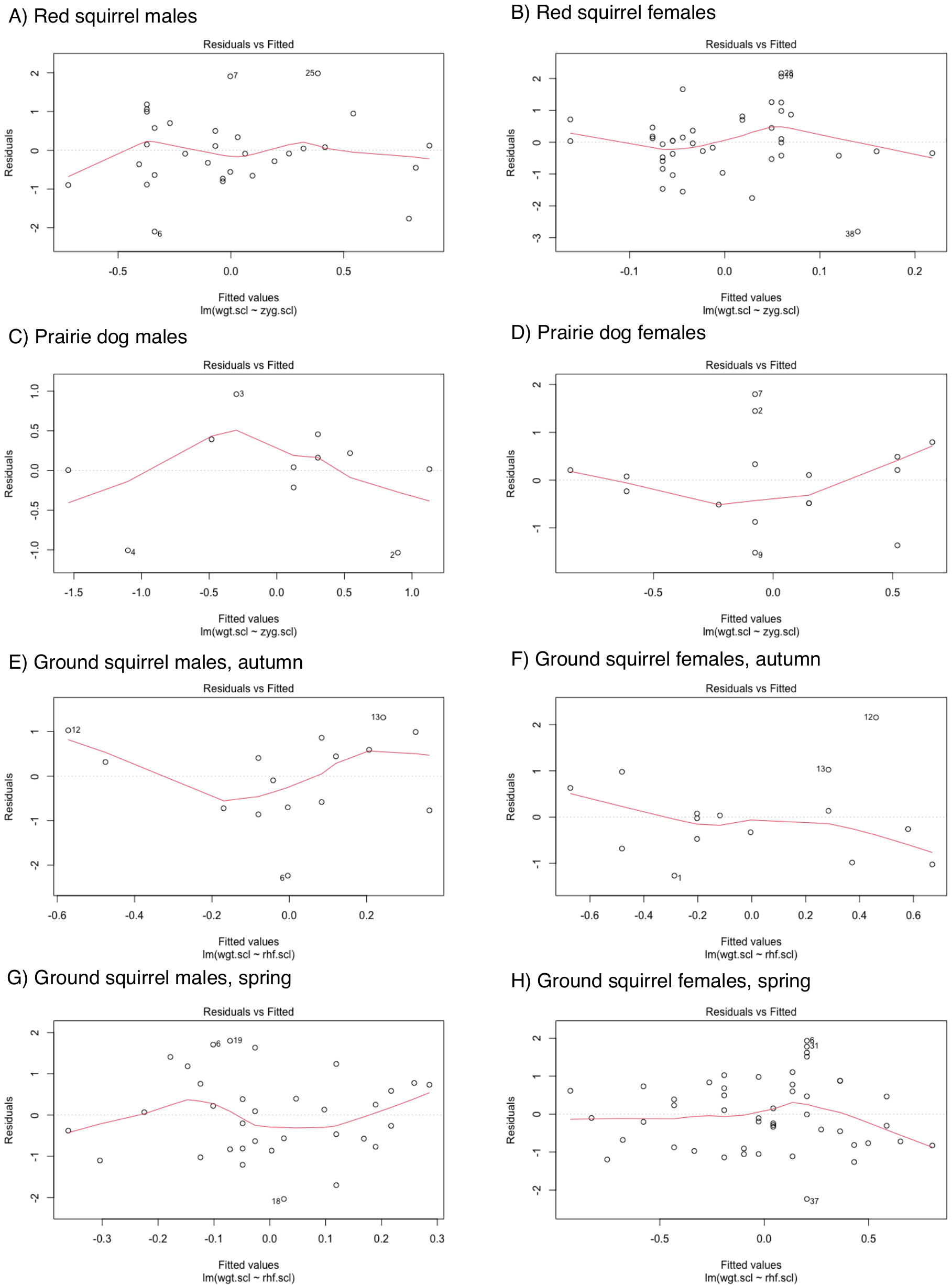

**Supplementary Figure S2.** Residual plots for a regression between body mass (log transformed and scaled within sex within species) and the skeletal measure used to derive the body condition index (either right hind foot length [RHF] or zygomatic width [ZYG; both log transformed and scaled within sex within species]) for A) North American red squirrel males pre-winter, B) red squirrel males pre-winter, C) black-tailed prairie dog males pre-winter, D) prairie dog females pre-winter, E) Columbian ground squirrel males pre-winter, F) ground squirrel females pre-winter, G) ground squirrel males in spring, and H) ground squirrel females in spring.

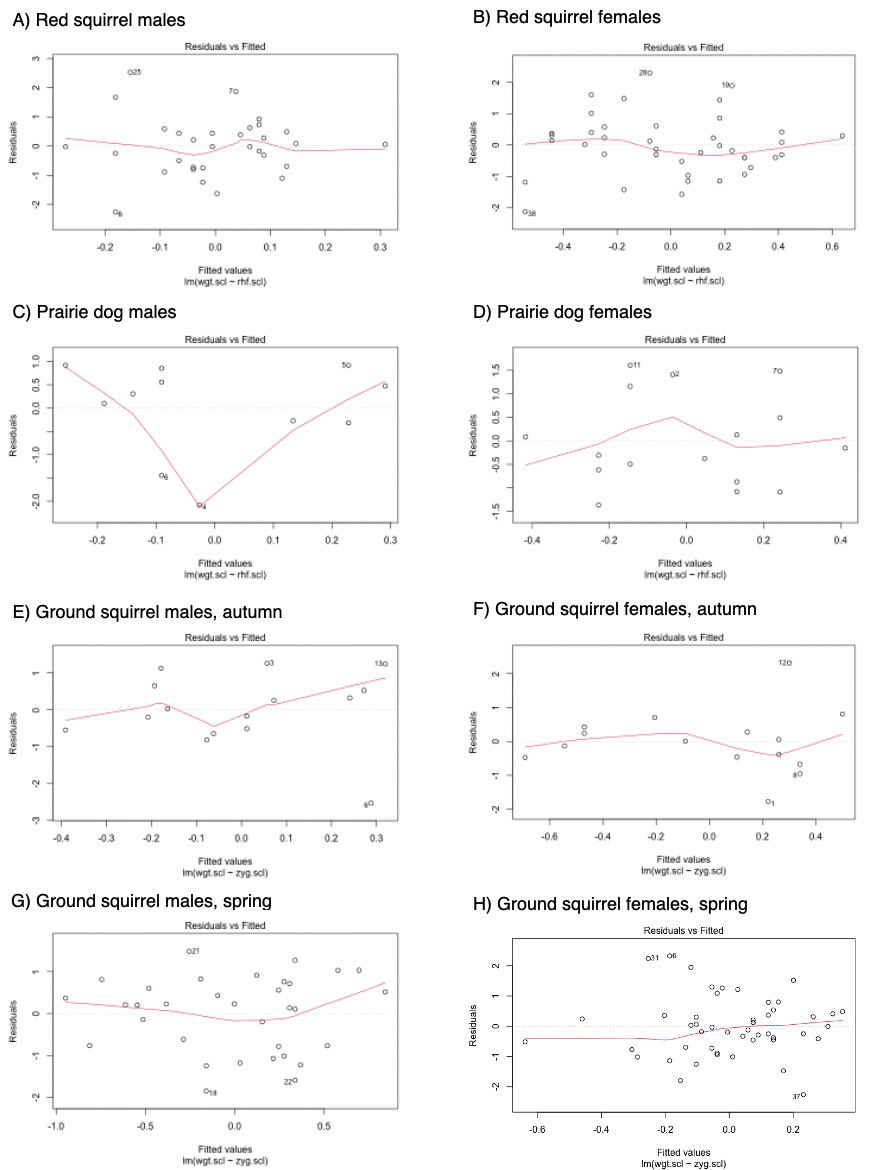

**Supplementary Figure S3.** Residual plots for a regression between body mass (log transformed and scaled within sex within species) and the alternate skeletal measure that was not used to derive the body condition index (either right hind foot length [RHF] or zygomatic width [ZYG; both log transformed and scaled within sex within species]) for A) North American red squirrel males pre-winter, B) red squirrel males pre-winter, C) black-tailed prairie dog males pre-winter, D) prairie dog females pre-winter, E) Columbian ground squirrel males pre-winter, F) ground squirrel females pre-winter, G) ground squirrel males in spring, and H) ground squirrel females in spring.

**Table S1.** Pearson product moment correlation coefficients between right hind foot length and zygomatic width for male and female North American red squirrels, black-tailed prairie dogs, and Columbian ground squirrels.

| **Species** | **Sex** | ***R*** | **p-value** |
| --- | --- | --- | --- |
| North American red squirrels | ♂ | -0.147 | 0.430 |
|  | ♀ | -0.154 | 0.342 |
| Black-tailed prairie dogs | ♂ | 0.553 | 0.078 |
|  | ♀ | -0.246 | 0.359 |
| Columbian ground squirrels  Spring | ♂ | 0.014 | 0.937 |
|  | ♀ | 0.118 | 0.425 |
| Pre-winter | ♂ | -0.306 | 0.267 |
|  | ♀ | 0.681 | 0.005* |
| **denotes significant correlation, Pearson's product moment correlation with α = 0.05* | | | |

**Table S2.** Coefficients of variation (CV) for body mass, morphometric measurements (right hind foot length = RHF, zygomatic width = ZW), and body composition (fat and lean mass) for male and female North American red squirrels, black-tailed prairie dogs, and Columbian ground squirrels. Bold values indicate whether RHF or ZW was used to calculate body condition index for each species.

|  |  | **Coeffecient of variation (CV)** | | | | |
| --- | --- | --- | --- | --- | --- | --- |
| **Species** | **Sex** | **Body mass (g)** | **RHF (mm)** | **ZW (mm)** | **Fat (g)** | **Lean (g)** |
| North American red squirrels | ♂ | 5.96 | 2.76 | **4.24** | 13.92 | 5.15 |
|  | ♀ | 7.09 | 3.08 | **2.78** | 14.08 | 8.77 |
| Black-tailed prairie dogs | ♂ | 6.23 | 2.06 | **2.67** | 14.06 | 6.53 |
|  | ♀ | 9.21 | 1.48 | **2.42** | 17.66 | 9.41 |
| Columbian ground squirrels  *Spring* | ♂ | 11.42 | **4.58** | 4.04 | 30.40 | 11.37 |
|  | ♀ | 12.27 | **3.75** | 3.70 | 49.05 | 11.87 |
| *Pre-winter* | *♂* | 6.23 | **4.43** | 4.05 | 14.06 | 6.53 |
|  | ♀ | 9.21 | **3.22** | 3.70 | 17.66 | 9.41 |

**Table S3.** Model coefficients for linear models testing whether a body condition index (BCI) derived from zygomatic width (ZW index) or body mass better explains observed variation in fat and lean mass in North American red squirrels pre-winter. Reference group for sex: female.

| **Component** | **Model** | **Independent terms** | **Estimate** | ***t*** | ***p*** | **AICc** | **Adjusted *R^2^*** |
| --- | --- | --- | --- | --- | --- | --- | --- |
| Fat | ~ BCI*sex | Intercept | 7.22 ± 0.44 | 16.54 | < 0.001 * | 390.7 | 0.22 |
|  |  | BCI | 1.87 ± 0.44 | 4.26 | < 0.001 * |  |  |
|  |  | Sex | -1.25 ± 0.68 | -1.85 | 0.069 |  |  |
|  |  | BCI*sex | -1.03 ± 0.73 | -1.42 | 0.161 |  |  |
|  | ~mass*sex | Intercept | -21.67 ± 6.36 | -3.41 | < 0.002 * | 389.4 | 0.22 |
|  |  | Body mass (g) | 0.12 ± 0.03 | 4.56 | < 0.001 * |  |  |
|  |  | Sex | 19.26 ± 10.72 | 1.80 | 0.077 |  |  |
|  |  | Mass*sex | -0.09 ± 0.04 | -1.977 | 0.051 |  |  |
| Lean | ~ BCI*sex | Intercept | 214.51 ± 1.33 | 160.89 | < 0.001 * | 562.6 | 0.64 |
|  |  | BCI | 12.28 ± 1.34 | 9.12 | < 0.001 * |  |  |
|  |  | Sex | 3.46 ± 2.07 | 1.67 | 0.099 |  |  |
|  |  | BCI*sex | 0.85 ± 2.22 | 0.38 | 0.702 |  |  |
|  | ~mass*sex | Intercept | 24.13 ± 15.53 | 1.55 | 0.125 | 527.0 | 0.78 |
|  |  | Body mass (g) | 0.78 ± 0.06 | 12.30 | < 0.001 * |  |  |
|  |  | Sex | -28.60 ± 26.20 | -1.09 | 0.279 |  |  |
|  |  | Mass*sex | 0.10 ± 0.11 | 1.00 | 0.323 |  |  |

**denotes significant p-value at α = 0.05*

**Table S4.** Model coefficients for linear models testing whether a body condition index (BCI) derived from zygomatic width (ZW index) or body mass better explains observed variation in fat and lean mass in black-tailed prairie dogs pre-winter. Reference group for sex: female.

| **Component** | **Model** | **Independent terms** | **Estimate** | ***t*** | ***p*** | **AICc** | **Adjusted *R^2^*** |
| --- | --- | --- | --- | --- | --- | --- | --- |
| Fat | ~ BCI*sex | Intercept | 517.47 ± 10.48 | 49.36 | < 0.001 * | 286.9 | 0.65 |
|  |  | BCI | 69.42 ± 12.04 | 5.77 | < 0.001 * |  |  |
|  |  | Sex | -21.91 ± 16.43 | -1.33 | 0.195 |  |  |
|  |  | BCI*sex | 20.57 ± 25.45 | 0.808 | 0.427 |  |  |
|  | ~mass*sex | Intercept | -397.6 ± 128.5 | -3.41 | 0.005 * | 277.4 | 0.75 |
|  |  | Body mass (g) | 0.628 ± 0.09 | 7.14 | < 0.001 * |  |  |
|  |  | Sex | -109.1 ± 227.0 | -0.48 | 0.635 |  |  |
|  |  | Mass*sex | -0.00 ± 0.15 | -0.00 | 0.999 |  |  |
| Lean | ~ BCI*sex | Intercept | 727.53 ± 12.38 | 58.79 | < 0.001 * | 295.9 | 0.82 |
|  |  | BCI | 47.44 ± 14.21 | 3.34 | 0.003 * |  |  |
|  |  | Sex | 201.26 ± 19.39 | 10.38 | < 0.001 * |  |  |
|  |  | BCI*sex | -56.48 ± 30.04 | -1.88 | 0.073 |  |  |
|  | ~mass*sex | Intercept | -56.00 ± 140.98 | -0.40 | 0.695 | 282.4 | 0.89 |
|  |  | Body mass (g) | 0.54 ± 0.10 | 5.57 | < 0.001 * |  |  |
|  |  | Sex | 675 ± 249.01 | 2.71 | 0.013 * |  |  |
|  |  | Mass*sex | -0.34 ± 0.16 | -2.14 | 0.043 |  |  |

**denotes significant p-value at α = 0.05*

**Table S5.** Model coefficients for linear models testing whether a body condition index (BCI) derived from right hind foot length (RHF index) or body mass better explains observed variation in fat and lean mass in Columbian ground squirrels pre-winter. Reference group for sex: female.

| **Component** | **Model** | **Independent terms** | **Estimate** | ***t*** | ***p*** | **AICc** | **Adjusted *R^2^*** |
| --- | --- | --- | --- | --- | --- | --- | --- |
| Fat | ~ BCI*sex | Intercept | 116.42 ± 4.53 | 25.71 | < 0.001 * | 265.2 | 0.80 |
|  |  | BCI | 15.68 ± 5.16 | 3.04 | 0.005 * |  |  |
|  |  | Sex | 62.85 ±6.40 | 9.81 | < 0.001 * |  |  |
|  |  | BCI*sex | 1.78 ± 7.09 | 0.25 | 0.804 |  |  |
|  | ~mass*sex | Intercept | -49.70 ± 52.39 | -0.95 | 0.352 | 266.7 | 0.80 |
|  |  | Body mass (g) | 0.30 ± 0.09 | 3.18 | 0.004 * |  |  |
|  |  | Sex | -10.43 ± 93.38 | -0.11 | 0.912 |  |  |
|  |  | Mass*sex | 0.04 ± 0.14 | 0.26 | 0.794 |  |  |
| Lean | ~ BCI*sex | Intercept | 388.37 ± 5.73 | 67.81 | < 0.001 * | 279.3 | 0.83 |
|  |  | BCI | 34.54 ± 6.52 | 5.30 | < 0.001 * |  |  |
|  |  | Sex | 83.35 ± 8.1 | 10.29 | < 0.001 * |  |  |
|  |  | BCI*sex | -14.23 ± 8.96 | -1.89 | 0.124 |  |  |
|  | ~mass*sex | Intercept | 12.61 ± 48.80 | 0.26 | 0.789 | 262.5 | 0.90 |
|  |  | Body mass (g) | 0.67 ± 0.09 | 7.73 | < 0.001 * |  |  |
|  |  | Sex | 78.36 ± 87.00 | 0.90 | 0.376 |  |  |
|  |  | Mass*sex | -0.14 ± 0.13 | -1.053 | 0.302 |  |  |

**denotes significant p-value at α = 0.05*

**Table S6.** Model coefficients for linear models testing whether a body condition index (BCI) derived from right hind foot length (RHF index) or body mass better explains observed variation in fat and lean mass in Columbian ground squirrels in spring. Reference group for sex: female.

| **Component** | **Model** | **Independent terms** | **Estimate** | ***t*** | ***p*** | **AICc** | **Adjusted *R^2^*** |
| --- | --- | --- | --- | --- | --- | --- | --- |
| Fat | ~ BCI*sex | Intercept | 29.50 ± 2.03 | 14.56 | < 0.001 * | 664.6 | 0.59 |
|  |  | BCI | 8.34 ± 2.23 | 3.74 | < 0.001 * |  |  |
|  |  | Sex | 30.16 ± 3.18 | 9.50 | < 0.001 * |  |  |
|  |  | BCI*sex | 0.50 ± 3.36 | 0.148 | 0.883 |  |  |
|  | ~mass*sex | Intercept | -35.48 ± 16.84 | -2.11 | 0.038 * | 664.9 | 0.59 |
|  |  | Body mass (g) | 0.17 ± 0.04 | 3.87 | < 0.001 * |  |  |
|  |  | Sex | 23.41 ± 27.65 | 0.85 | 0.400 |  |  |
|  |  | Mass*sex | -0.02 ± 0.06 | -0.35 | 0.729 |  |  |
| Lean | ~ BCI*sex | Intercept | 310.09 ± 2.55 | 121.85 | < 0.001 * | 701.5 | 0.89 |
|  |  | BCI | 34.54 ± 6.52 | 12.41 | < 0.001 * |  |  |
|  |  | Sex | 70.46 ± 3.99 | 17.67 | < 0.001 * |  |  |
|  |  | BCI*sex | 5.95 ± 4.22 | 1.41 | 0.163 |  |  |
|  | ~mass*sex | Intercept | 22.57 ± 14.30 | 1.58 | 0.119 | 638.3 | 0.95 |
|  |  | Body mass (g) | 0.76 ± 0.04 | 20.25 | < 0.001 * |  |  |
|  |  | Sex | -2.22 ± 23.47 | -0.10 | 0.925 |  |  |
|  |  | Mass*sex | -0.001 ± 0.05 | -0.15 | 0.880 |  |  |

**denotes significant p-value at α = 0.05*

**Table S7.** Model coefficients for linear models testing whether a body condition index (BCI) derived from right hind foot length (RHF index) explains observed variation in fat and lean mass in North American red squirrels pre-winter. Compare to Supplementary Table S3.

| **Component** | **Model** | **Independent terms** | **Estimate** | ***t*** | ***p*** | **AICc** | **Adjusted *R^2^*** |
| --- | --- | --- | --- | --- | --- | --- | --- |
| Fat | ~ BCI*sex | Intercept | 7.17 ± 0.46 | 15.67 | < 0.001 * | 397.8 | 0.139 |
|  |  | BCI | 1.67 ± 0.49 | 3.41 | 0.001 * |  |  |
|  |  | Sex (ref: male) | -1.22 ± 0.71 | -1.713 | 0.091 |  |  |
|  |  | BCI*sex (male) | -1.35± 0.74 | -1.81 | 0.074 |  |  |
| Lean | ~ BCI*sex | Intercept | 214.12 ± 1.07 | 200.803 | < 0.001 * | 528.1 | 0.773 |
|  |  | BCI | 13.40 ± 1.14 | 11.76 | < 0.001 * |  |  |
|  |  | Sex (ref: male) | 3.88 ± 1.65 | 2.35 | < 0.001 * |  |  |
|  |  | BCI*sex (male) | 0.91 ± 1.73 | 0.52 | 0.603 |  |  |

**denotes significant p-value at α = 0.05*

**Table S8.** Model coefficients for linear models testing whether a body condition index (BCI) derived from right hind foot length (RHF index) explains observed variation in fat and lean mass in black-tailed prairie dogs pre-winter. Compare to Supplementary Table S4.

| **Component** | **Model** | **Independent terms** | **Estimate** | ***t*** | ***p*** | **AICc** | **Adjusted *R^2^*** |
| --- | --- | --- | --- | --- | --- | --- | --- |
| Fat | ~ BCI*sex | Intercept | 517.47 ± 7.89 | 65.61 | < 0.001 * | 271.5 | 0.8015 |
|  |  | BCI | 68.01 ± 8.37 | 8.13 | 0.001 * |  |  |
|  |  | Sex (ref: male) | -21.91 ± 12.36 | -1.77 | 0.090 |  |  |
|  |  | BCI*sex (male) | -4.91 ± 13.16 | -0.37 | 0.712 |  |  |
| Lean | ~ BCI*sex | Intercept | 727.53 ± 10.94 | 66.48 | < 0.001 * | 289.2 | 0.857 |
|  |  | BCI | 51.76 ± 11.61 | 4.46 | 0.002 * |  |  |
|  |  | Sex (ref: male) | 201.26 ± 17.14 | 11.74 | < 0.001 * |  |  |
|  |  | BCI*sex (male) | -38.04 ± 18.26 | -2.08 | 0.049 * |  |  |

**denotes significant p-value at α = 0.05*

**Table S9.** Model coefficients for linear models testing whether a body condition index (BCI) derived from zygomatic width (ZW index) explains observed variation in fat and lean mass in Columbian ground squirrels pre-winter. Compare to Supplementary Table S5.

| **Component** | **Model** | **Independent terms** | **Estimate** | ***t*** | ***p*** | **AICc** | **Adjusted *R^2^*** |
| --- | --- | --- | --- | --- | --- | --- | --- |
| Fat | ~ BCI*sex | Intercept | 116.42 ± 4.69 | 24.83 | < 0.001 * | 267.3 | 0.485 |
|  |  | BCI | 14.44 ± 5.25 | 2.75 | 0.011 * |  |  |
|  |  | Sex (ref: male) | 62.85 ± 6.63 | 9.48 | < 0.001 * |  |  |
|  |  | BCI*sex (male) | 2.31 ± 7.23 | 0.32 | 0.752 |  |  |
| Lean | ~ BCI*sex | Intercept | 388.37 ± 5.42 | 71.69 | < 0.001 * | 276.0 | 0.848 |
|  |  | BCI | 34.17 ± 6.07 | 5.63 | < 0.001 * |  |  |
|  |  | Sex (ref: male) | 83.35 ± 7.66 | 10.88 | < 0.001 * |  |  |
|  |  | BCI*sex (male) | -12.00 ± 8.354 | -1.44 | 0.163 |  |  |

**denotes significant p-value at α = 0.05*

**Table S10.** Model coefficients for linear models testing whether a body condition index (BCI) derived from zygomatic width (ZW index) explains observed variation in fat and lean mass in Columbian ground squirrels in spring. Compare to Supplementary Table S6.

| **Component** | **Model** | **Independent terms** | **Estimate** | ***t*** | ***p*** | **AICc** | **Adjusted *R^2^*** |
| --- | --- | --- | --- | --- | --- | --- | --- |
| Fat | ~ BCI*sex | Intercept | 29.51 ± 2.03 | 14.55 | < 0.001 * | 664.7 | 0.586 |
|  |  | BCI | 7.79 ± 2.01 | 3.72 | < 0.001 * |  |  |
|  |  | Sex (ref: male) | 30.16 ± 3.18 | 3.18 | < 0.001 * |  |  |
|  |  | BCI*sex (male) | 1.97± 3.48 | 0.57 | 0.573 |  |  |
| Lean | ~ BCI*sex | Intercept | 310.09 ± 2.74 | 113.25 | < 0.001 * | 713.3 | 0.870 |
|  |  | BCI | 35.47 ± 2.83 | 12.56 | < 0.001 * |  |  |
|  |  | Sex (ref: male) | 70.46 ± 4.29 | 16.42 | < 0.001 * |  |  |
|  |  | BCI*sex (male) | 3.69 ± 4.70 |  | 0.435 |  |  |

**denotes significant p-value at α = 0.05*

**Table S11.** Comparing candidate models for predicting fat mass and lean mass from either scaled mass index (SMI) or body mass (scaled within species) within species and season for North American red squirrels (pre-winter), black-tailed prairie dogs p(re-winter), and Columbian ground squirrels (pre-winter and spring). Akaike information criterion (corrected for small sample sizes; AICc) and adjusted *R^2^* values for all mass*sex models are the same as those presented for identical models shown in Supplementary Tables S3-S6.

| **Species/Season** | **Response variable (body component)** | **Model** | **AICc** | **Adjusted *R^2^*** |
| --- | --- | --- | --- | --- |
| Red squirrels,  pre-winter | Fat | ~ SMI*sex | 386.8 | 0.25 |
|  |  | ~mass*sex | 389.4 | 0.22 |
|  | Lean | ~ SMI*sex | 626.4 | 0.18 |
|  |  | ~mass*sex | 527.0 | 0.78 |
| Prairie dogs,  pre-winter | Fat | ~ SMI *sex | 306.9 | 0.26 |
|  |  | ~mass*sex | 277.4 | 0.75 |
|  | Lean | ~ SMI*sex | 304.9 | 0.74 |
|  |  | ~mass*sex | 282.4 | 0.89 |
| Ground squirrels,  pre-winter | Fat | ~ SMI*sex | 271.7 | 0.75 |
|  |  | ~mass*sex | 266.7 | 0.80 |
|  | Lean | ~ SMI*sex | 301.4 | 0.65 |
|  |  | ~mass*sex | 262.5 | 0.90 |
| Ground squirrels,  spring | Fat | ~ SMI*sex | 677.3 | 0.52 |
|  |  | ~mass*sex | 664.9 | 0.59 |
|  | Lean | ~ SMI*sex | 802.8 | 0.64 |
|  |  | ~mass*sex | 638.3 | 0.95 |

**Table S12.** Model coefficients for a set of linear models testing the interactive and singular effects of body condition index (BCI) or body mass (scaled within species) and species on fat and lean mass in red squirrels, prairie dogs, and ground squirrels (pre-winter and spring). Reference species: red squirrels.

| **Predictor** | **Component** | **Independent terms** | **Estimate** | ***t*** | ***p*** | **AICc** | **Adjusted *R^2^*** |
| --- | --- | --- | --- | --- | --- | --- | --- |
| BCI | Fat (g) | Intercept | 6.70 ± 2.70 | 2.485 | 0.014 * | 1981.6 | 0.98 |
|  |  | BCI | 1.57 ± 2.84 | 0.53 | 0.596 |  |  |
|  |  | Species (pre-winter prairie dogs) | 501.84 ± 5.30 | -94.78 | < 0.001 * |  |  |
|  |  | Species (pre-winter ground squirrels) | -141.14 ± 5.10 | 27.70 | < 0.001 * |  |  |
|  |  | Species (spring ground squirrels) | 35.09 ± 3.77 | 9.31 | < 0.001 * |  |  |
|  |  | BCI*species (pre-winter prairie dogs) | 72.52 ± 6.63 | 10.94 | < 0.001 * |  |  |
|  |  | BCI*species (pre-winter ground squirrels) | 15.11 ± 5.55 | 2.73 | 0.007 * |  |  |
|  |  | BCI*species (spring ground squirrels) | 7.05 ± 4.00 | 1.77 | 0.079 |  |  |
|  | Lean (g) | Intercept | 215.94 ± 5.74 | 37.70 | < 0.001 * | 2305.3 | 0.93 |
|  |  | BCI | 12.56 ± 6.02 | 2.09 | 0.039 * |  |  |
|  |  | Species (pre-winter prairie dogs) | 593.58 ± 11.24 | 52.80 | < 0.001 * |  |  |
|  |  | Species (pre-winter ground squirrels) | 214.10 ± 10.82 | 19.79 | < 0.001 * |  |  |
|  |  | Species (spring ground squirrels) | 122.85 ± 8.00 | 15.36 | < 0.001 * |  |  |
|  |  | BCI*species (pre-winter prairie dogs) | 22.24 ± 14.07 | 1.58 | 0.115 |  |  |
|  |  | BCI*species (pre-winter ground squirrels) | 14.43 ± 11.79 | 1.23 | 0.222 |  |  |
|  |  | BCI*species (spring ground squirrels) | 24.84 ± 8.48 | 2.93 | 0.004 * |  |  |
| Body mass (scaled) | Fat (g) | Intercept | 6.69 ± 2.54 | 2.64 | < 0.001 * | 1954.9 | 0.98 |
|  |  | Body mass (scaled) | 1.40 ± 2.62 | 0.54 | < 0.001 * |  |  |
|  |  | Species (pre-winter prairie dogs) | 501.85 ± 54.98 | 100.84 | < 0.001 * |  |  |
|  |  | Species (pre-winter ground squirrels) | 141.15 ± 4.79 | 29.47 | < 0.001 * |  |  |
|  |  | Species (spring ground squirrels) | 35.10 ± 3.54 | 9.91 | < 0.001 * |  |  |
|  |  | Mass *species (prairie dogs) | 61.61 ± 65.16 | 11.93 | < 0.001 * |  |  |
|  |  | Mass*species (pre-winter ground squirrels) | 13.74 ± 4.95 | 2.77 | 0.006 * |  |  |
|  |  | Mass *species (spring ground squirrels) | 6.92 ± 3.62 | 1.91 | 0.057 |  |  |
|  | Lean (g) | Intercept | 215.83 ± 5.49 | 39.29 | < 0.001 * | 2287.3 | 0.94 |
|  |  | Body mass (scaled) | 13.42 ± 5.67 | 2.37 | 0.019 |  |  |
|  |  | Species (pre-winter prairie dogs) | 593.69 ± 10.78 | 55.06 | < 0.001 * |  |  |
|  |  | Species (pre-winter ground squirrels) | 214.21 ± 10.38 | 20.65 | < 0.001 * |  |  |
|  |  | Species (spring ground squirrels) | 122.96 ± 7.67 | 16.03 | < 0.001 * |  |  |
|  |  | Mass *species (pre-winter prairie dogs) | 27.89 ± 11.18 | 2.47 | 0.014 * |  |  |
|  |  | Mass *species (pre-winter ground squirrels) | 15.67 ± 10.73 | 1.46 | 0.146 |  |  |
|  |  | Mass *species (spring ground squirrels) | 23.89 ± 7.85 | 3.05 | 0.003 * |  |  |

**denotes significant p-value at α = 0.05*
